# Inductive determination of allele frequency spectrum probabilities in structured populations

**DOI:** 10.1101/454157

**Authors:** Marcy K. Uyenoyama, Naoki Takebayashi, Seiji Kumagai

## Abstract

We present a method for inductively determining exact allele frequency spectrum (AFS) probabilities for samples derived from a population comprising two demes under the infinite-allele model of mutation. This method builds on a labeled coalescent argument to extend the Ewens sampling formula (ESF) to structured populations. A key departure from the panmictic case is that the AFS conditioned on the number of alleles in the sample is no longer independent of the scaled mutation rate (*θ*). In particular, biallelic site frequency spectra, widely-used in explorations of genome-wide patterns of variation, depend on the mutation rate in structured populations. Variation in the rate of substitution across loci and through time may contribute to apparent distortions of site frequency spectra exhibited by samples derived from structured populations.

---

Ewens (1972) addressed the sampling distribution of genes derived from a panmictic population comprising an effective number of 2*N* autosomal genes under the infinite-allele model of mutation. Novel alleles are generated at the per-gene, per-generation rate of *u*, with

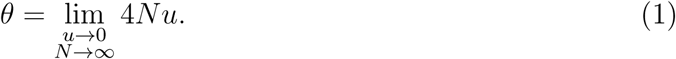

A sample comprising *n* genes has allele frequency spectrum

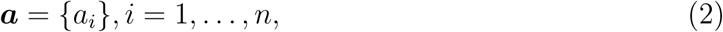

for *a_i_* the number of alleles that occur with multiplicity *i* in the sample. The Ewens Sampling Formula (ESF) provides the probability of the sample:

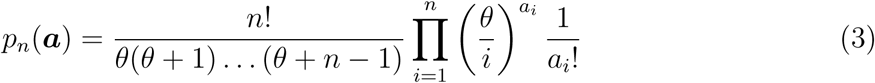

Ewens 1972; Karlin and McGregor 1972).

A remarkable property of the ESF is that the probability that the next-sampled gene represents a novel allelic class depends only on the size of the sample and the mutation rate. This property constitutes a fundamental link between the ESF and an extensive range of stochastic structures (Tavaré and Ewens 1997), including urn models (Hoppe 1987), the Chinese Restaurant Process (Aldous 1985), and the coalescent itself (Kingman 2000). A novel allele is observed on the *n^th^* draw with probability

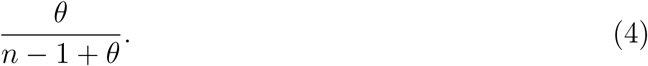

Alternatively, the *n^th^* gene belongs to an allelic class already represented by *i* replicates with probability

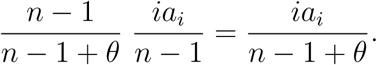

While the probability that the next-sampled gene is novel (4) is well-known, the usual demonstration involves observing that it is a corollary of the ESF (*e.g.*, Karlin and McGregor 1972). Redelings *et al.* (2015) presented a first-principles derivation of (4) that does not require knowledge of the ESF in advance.

Another remarkable property of the ESF is that the number of alleles observed in a sample (*K*) provides a sufficient statistic for the estimation of the scaled mutation rate *θ*, the sole parameter of the ESF (Ewens 1972): conditioning the ESF on a particular value of *K* results in cancellation of *θ*. Ganapathy and Uyenoyama (2009) observed that for arbitrary values of *θ*, the ESF conditioned on *K* = 2 alleles corresponds to the folded site frequency spectrum (SFS). This independence of the SFS from the mutation rate at each site has proven especially useful in genome-wide analyses of biallelic SNP variation, obviating the need to account for variation in mutation rates across regions or to assume vanishingly small mutation rates at SNP loci.

Here, we present an inductive method for generating the exact probability mass function of the allele frequency spectrum (AFS) in a sample of arbitrary size derived from a population comprising multiple demes. We begin with an overview of our approach, emphasizing key features introduced by population structure. We provide a Mathematica implementation of our method as a supplementary file.

## 1 Overview

### 1.1 Transition rates

We consider a two-deme model, in which deme *i* (*i* = 0, 1) comprises an effective number of 2*N_i_* genes with backward migration rate *m_i_*. Let *r* denote the proportion of reproductive individuals that reside in deme 0:

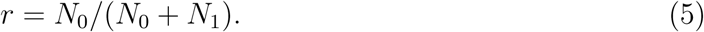

As in the ESF, mutation follows the infinite-allele model, with novel alleles generated at the per-gene, per-generation rate of *u*. We consider the rare-event, large-population case, for which

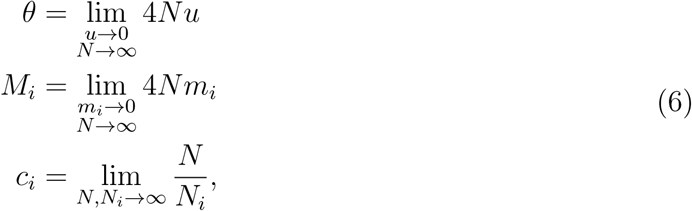

in which *N* represents an arbitrary quantity that goes to infinity at the same rate as *N*_0_ and *N*_1_.

While our model has 5 basic parameters (*u*, *m*_0_, *m*_1_, *c*_0_, *c*_1_), the coalescence approach makes clear that the pattern of variation in the sample depends on only relative rates, implying just 4 degrees of freedom. For example, assigning

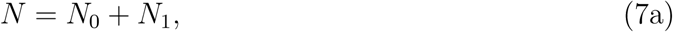

implies

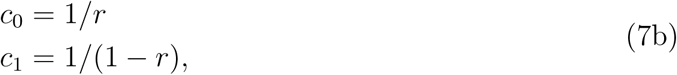

for *r* given in (5). Alternatively, assigning *N* as the unweighted mean of effective numbers,

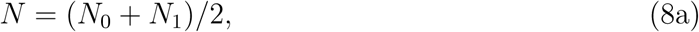

implies

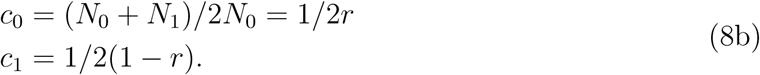

To promote clarity and facilitate comparison to previous work, we assume at points in our exposition complete symmetry:

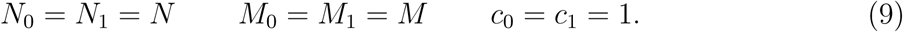

We address the probability of allele frequency spectrum (AFS) ***a*** observed in a sample comprising *n_i_* genes from deme *i* (*i* = 0, 1):

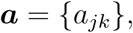

for *a_jk_* the number of alleles with exactly *j* replicates in deme 0 and *k* in deme 1. Let *p*_*n*_0_,*n*_1__ (***a***) denote the probability of observing this AFS. We use the framework described by Kumagai and Uyenoyama (2015) to develop the central recursion in AFS probabilities in this two-deme setting (Appendix A). Expansion of the method to accommodate more than two demes would entail a larger linear system (Section 4.1), but would preserve the essence of the coalescence-based approach to labeled samples.

### 1.2 Coalescence of labeled lineages

Karlin and McGregor’s (1972) recursion in the allele frequency spectrum constitutes a *labeled* coalescent argument, addressing the descent of lineages of specified allelic types. Observation of allelic class clearly excludes certain events in the immediate past (*e.g.*, coalescence of members of distinct allelic classes). However, unexcluded events are not in general equally likely, with the observed pattern of variation providing information about the history of the sample (see Wiuf and Donnelly 1999).

Because exhaustive evaluation of the typically enormous number of possible ancestral states is impractical, importance sampling has figured prominently in coalescence-based likelihood methods (Griffiths and Tavaré 1994; Felsenstein *et al.* 1999; Stephens and Donnelly 2000). In a number of important works, De Iorio and Griffiths and colleagues (*e.g.*, De Iorio and Griffiths 2004; De Iorio *et al.* 2005) have developed importance sampling methods for the approximation of likelihoods under generalized mutation models for subdivided population schemes similar to ours.

Central to the efficiency of the importance sampling proposals of De Iorio and Griffiths and colleagues is the approximation of the probability that the next-sampled gene represents a novel allelic class. In structured populations, the allelic class of the last-sampled gene depends not only on sample size and *θ*, but also on other characteristics of the AFS of the present sample, including the number of genes sampled from each deme and the deme from which the last gene is derived.

The recursion in the AFS derived by Karlin and McGregor (1972) to prove the ESF constitutes a labeled coalescent argument rather than an importance sampling argument. Section 3.1 presents an exposition of the labeled coalescent recursion, extended to accommodate population structure. Previous works (*e.g.*, De Iorio and Griffiths 2004) have used similar arguments, but largely without comment. Our restriction to the infinite-allele model of mutation, as in the ESF for panmictic populations, permits inductive determination of the AFS probabilities without resort to importance sampling.

### 1.3 Conditioned and unconditioned frequency spectra

Using that the probability that the next gene sampled from a panmictic population represents a novel allele (4), Ewens (1972) derived the probability mass function of the number of alleles (*K*) observed in a sample of a given size. He showed that conditioning the ESF on the number of alleles produces a distribution that is independent of the scaled mutation rate (*θ*), the only parameter of the ESF.

Extending the approach described by Ganapathy and Uyenoyama (2009) to structured populations, we derive the probability generating function (pgf) for *K* (Section 2). This pgf provides the probability of a monomorphic sample (*K* = 1), the initial state for our inductive determination of the allele frequency spectrum (AFS). Further, conditioning the AFS probabilities on the observation of exactly two alleles produces the probability mass function (not only expectations) for the folded site frequency spectrum (SFS) for biallelic samples (Section 5.3). In structured populations, the number of alleles in the sample is no longer a sufficient statistic for the estimation of the scaled mutation rate: the SFS depends on all parameters (6), including *θ*.

## 2 Number of alleles in the sample

We use an *unlabeled* structured coalescent argument to derive the probability generating function (pgf) for *K*_*n*_0_,*n*_1__, the number of alleles observed in a sample comprising *n*_0_ genes derived from deme 0 and *n*_1_ from deme 1.

Termination of a lineage in a coalescence event establishes that it shares its allelic class with another lineage in the sample and, under the infinite-allele model of mutation, termination in mutation event implies that it represents a novel allelic class. Accordingly, to determine the number of alleles observed, we need follow a lineage up to only the most recent coalescence or mutation event (compare Griffiths and Lessard 2005). Ganapathy and Uyenoyama (2009) used this approach to generate the pgf for the panmictic case, an alternative path to the probability mass function derived by Ewens (1972).

Level ℓ of the gene genealogy corresponds to the segment of the genealogy in which exactly ℓ lineages exist. We index the states on level ℓ according to the number of lineages residing in deme 0. On level ℓ, the total rate of change from state *i* (*i* = 0, 1, …,ℓ) corresponds to

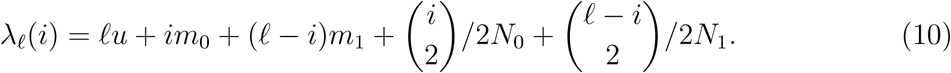

The Markov chain for each level terminates in either a mutation event or a coalescence event. In addition, the process may undergo transitions between transient states, reflecting migration events. For level ℓ of the gene genealogy, matrix ***U_ℓ_*** provides rates of transition among transient states. All elements of ***U_ℓ_*** are zero with the exception of those along the diagonal, superdiagonal, and subdiagonal, in accordance with the rare-event assumption (6), which restricts each migration event to a single lineage. For ***U_ℓ_***(*ij*) the element in row *i* and column *j* (*i, j* = 0, 1, …, ℓ),

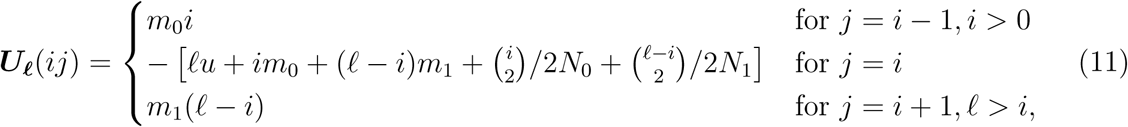

in which terms containing unmeaningful expressions 
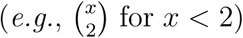
 are defined as zero. Rescaling using (10) as indicated in (A.7), we convert this matrix of transition rates into a matrix of transition probabilities (***Ũ***_***ℓ***_).

Matrix ***V_ℓ_*** provides rates of transition from transient states on level ℓ to absorption into level (ℓ − 1):

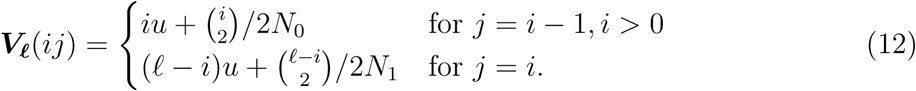

As for ***Ũ***_***ℓ***_, we use (10) and (A.7) to obtain ***V*̃**_***ℓ***_, a matrix of transition probabilities. As in (A.9), the element in the *i^th^* row and *j^th^* column of

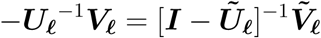

denotes the probability that a process initiated in state *i* of level ℓ is absorbed in state *j* of level (ℓ − 1).

In the numerator of

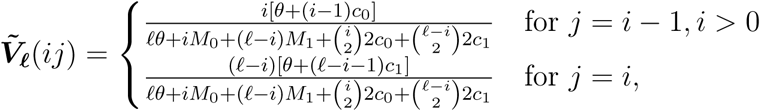

the *θ* indicates termination of level ℓ by mutation, implying the introduction of a new allelic class into the sample; otherwise, the level terminates through coalescence. To obtain the probability generating function for the number of alleles generated on level ℓ, we multiply each *θ* in the numerator of each element of ***V*̃**_***ℓ***_ by the parameter of the pgfs (*z*) to obtain

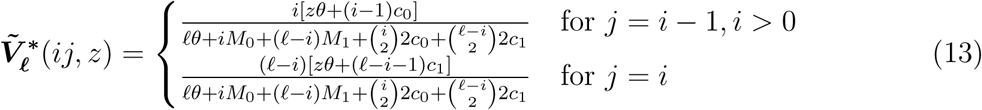

compare Kumagai and Uyenoyama 2015).

For each state of a sample comprising *n* genes, we construct a probability generating function that counts alleles in the sample, with lineages tracked backward in time until termination in either a mutation or coalescence event. The *i* + 1*^st^* element of ***G_n_***(*z*) gives the pgf for a sample in state *i* (*i* = 0, 1, …, *n*), indicating *i* lineages residing in deme 0. In general, ***G_n_***(*z*) is a vector of polynomials of degree (*n* − 1) in *z*, the parameter of the pgfs. Because ***G_n_***(*z*) counts the number of origins of a new allele, the probability of observing *K_i,n−i_* = *k* (*k* = 1, 2, …, *n*) alleles corresponds to the coefficient of *z*^(*k*−1)^ in the *i* + 1*^st^* element of ***G_n_***(*z*). The probabilities of monomorphic samples, comprising a single allele, are given by ***G_n_***(0).

For a sample of size 2,

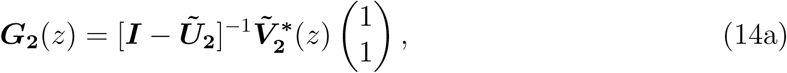

in which the vector on the right sums over the location (deme 0 or deme 1) of the most recent common ancestor. Starting with this base case, we obtain the pgfs for samples of successively larger size (ℓ > 2) by iterating

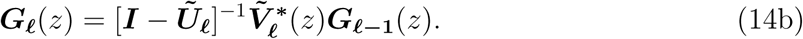

We obtain the probability of observing a monomorphic sample, with all genes representing the same allele, by setting *z* = 0 in the probability generating functions (14), implying termination of all levels of the gene genealogy by coalescence and not mutation. Under (8), the vector of probabilities of a monomorphic sample corresponds to

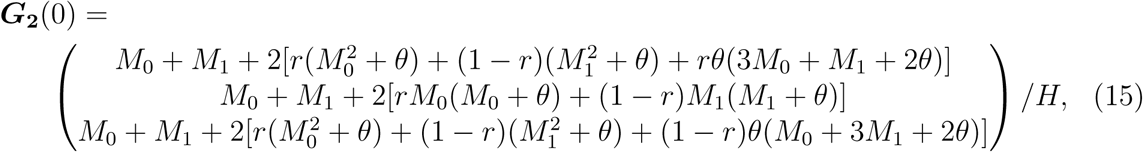

for

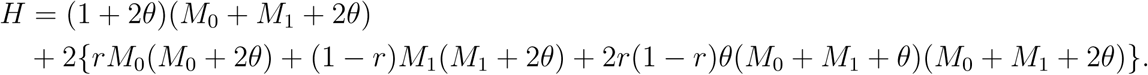

Under symmetry (9), which corresponds to *M*_0_ = *M*_1_ = *M* and *r* = 1/2 in (15), these expressions confirm the results of Hudson (1990). At the low-migration limit (*M*_0_, *M*_1_ → 0), corresponding to isolated demes, (14) gives

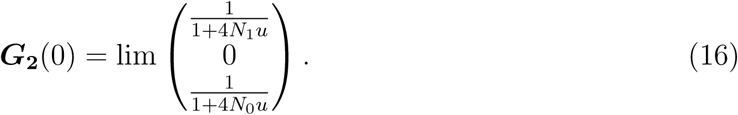

Under the infinite-allele model and in the absence of migration, genes sampled from distinct demes cannot belong to the same allelic class and genes sampled from the same deme are identical in state with probability

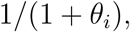

with scaled mutation rate appropriate for the local effective number (*θ_i_* = lim 4*N_i_u*). At the high-migration limit (*M*_0_, *M*_1_ → ∞), all elements of ***G*_2_**(0) reduce to

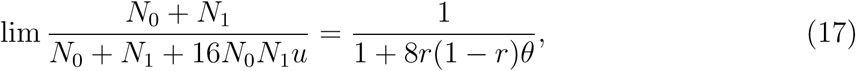

under (8), reflecting that the probability of monomorphism is independent of the original sampling configuration.

## 3 The labeled coalescent

Let random variable *D* represent the present (descendant) state of the sample and *A* the ancestral state, in which the most recent event (coalescence, mutation, or migration) occurs. We use a vertical argument (across time periods) to obtain a recursion relating *D* to *A*. A horizontal argument, which can reduce the computational load, appears in Section 4.2.

### 3.1 Vertical argument

For ***a*** the observed AFS, the likelihood Pr(*D* = ***a* Φ**) depends on the parameters of the model (6):

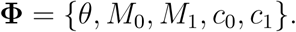

Random variable *T* indicates the event (coalescence, mutation, or migration) that joins descendant *D* to ancestor *A*. To determine Pr(*D* = ***a* Φ**), we consider all possible events and all possible ancestors:

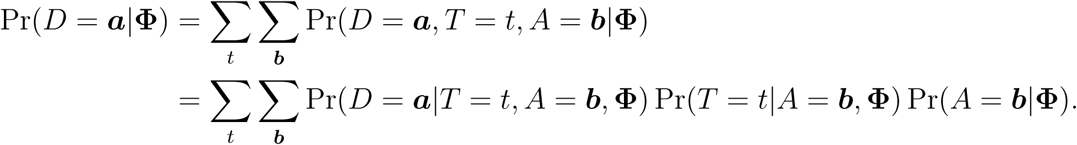

Viewed backward in time from descendant *D*, the most recent event is independent of the ancestor state:

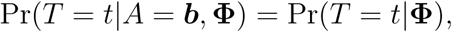

which implies

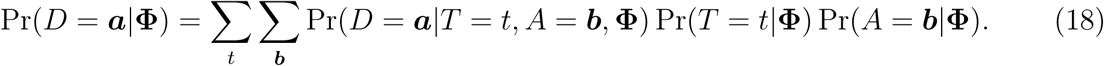

### 3.2 Transitions

We provide explicit expressions for the transitions that link the descendant probabilities Pr(*D|***Φ**) to the ancestor probabilities Pr(*A|***Φ**) in (18).

Observed allelic spectrum *D* = ***a*** comprises *n*_0_ genes in deme 0 and *n*_1_ in deme 1, for a total sample size of *n* = *n*_0_ + *n*_1_. The total per-generation rate of transition, through mutation, migration, or coalescence, corresponds to

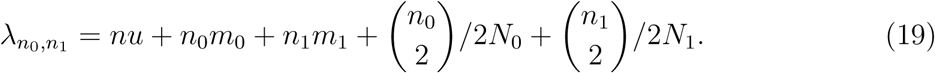

Event *T* may correspond to coalescence (*t*_0*i*_), mutation (*t*_1*i*_), or backward migration (*t*_2*i*_) in deme *i* (*i* = 0, 1). Coalescence reduces the number of lineages representing an allele. Mutation can occur only in a singleton (the sole representative of an allelic class) and induces no change in the number of lineages (*n*). Migration changes the partition of lineages among demes.

#### 3.2.1 Coalescence events

We first address formation of ancestor *A* by a coalescence event (*t*_00_ or *t*_01_). Under neutrality, ancestral events are independent on the allelic types of the lineages; accordingly, we determine the probabilities of coalescence events for *unlabeled* lineages:

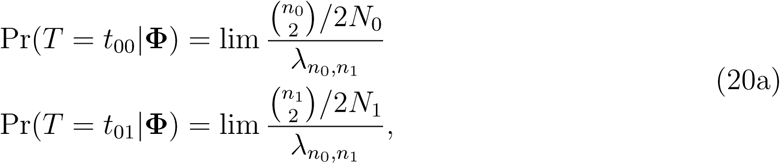

for *λ*_*n*0,*n*_1__ (19) the total rate of transition backward from *D* = ***a***. Some events may be incompatible with the descendant *D* = ***a***: for example, coalescence in deme 0 in which no allele with multiplicity greater than one resides. Such cases imply an ancestor *A* that has zero probability of generating descendant *D* through event *T*:

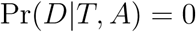

in (18).

For the event of coalescence in deme 0, any ancestor *A* = ***b*** with a positive probability of generating *D* = ***a*** must have one fewer allele relative to ***a*** with *i* replicates in deme 0 and *j* in deme 1 and one additional allele with *i* − 1 replicates in deme 0 and *j* in deme 1. Analogous considerations apply to the event of coalescence in deme 1. These constraints determine the compatible ancestors:

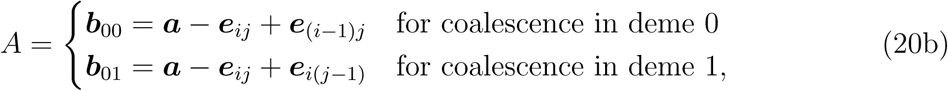

in which ***e_kl_*** represents a unit vector with the *kl* element corresponding to 1 and all other elements to 0.

Forward in time, a coalescence event entails the duplication of a lineage. Under the neutrality assumption, all lineages have identical probabilities of duplication, irrespective of allelic state. The probabilities of generating descendant *D* = ***a*** from compatible ancestors *A* (20b) through event *T* correspond to

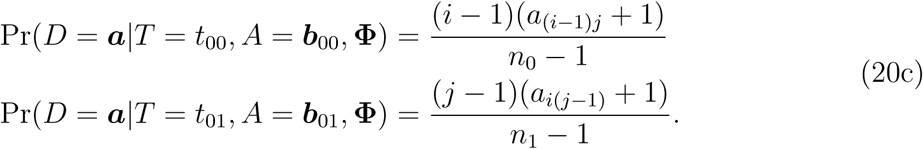

For example, event *T* = *t*_00_ entails the duplication of one of the genes in deme 0 of *A* = ***b***_00_ = ***a − e*** + ***e***_(*i*−1)*j*_. Of the (*n*_0_ − 1) genes in deme 0, (*a*_(*i*−1)*j*_ + 1) have (*i* − 1) replicates in deme 0 and *j* replicates in deme 1. Accordingly, the probability of forming *D* = ***a*** from *A* = ***b***_00_ through duplication of a gene in deme 0 corresponds to [(*i* − 1)*a*_(*i*−1)*j*_ + 1]/(*n*_0_ *−* 1).

#### 3.2.2 Mutation events

We now address formation of ancestor *A* by a mutation event (*t*_10_ or *t*_11_). Irrespective of the allelic types of the lineages, the probabilities of these events correspond to

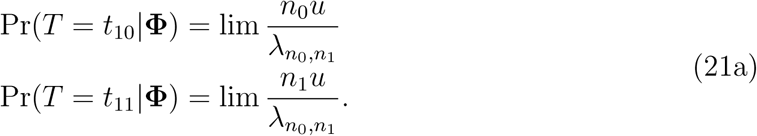

Under the infinite-allele model, mutation generates only novel alleles. Backward in time, mutation only in those lineages that represent a singleton allele in *D* can form an ancestor *A* with positive probability of generating *D*. Forward in time, the mutation may occur in a lineage that represents a singleton in *A*, in which case the allele frequency spectra of ancestor and descendant are identical (*A* = ***a***). Alternatively, the mutation may occur in a lineage of *A* that represents a non-singleton allele. For a mutation in deme 0, ancestor spectra with positive probability of giving rise to descendant *D* = ***a*** correspond to

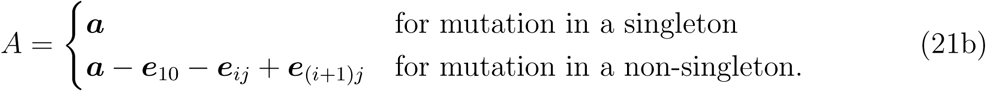

Similarly, the possible ancestor spectra for a mutation in deme 1 correspond to

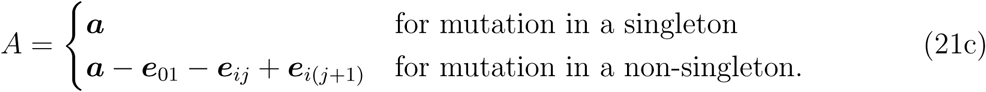

Note that the mutation event generates two singletons under the assignments (*i* = 1, *j* = 0) or (*i* = 0, *j* = 1).

The probabilities of generating descendant *D* = ***a*** from these ancestors *A* through the specified mutation event correspond to

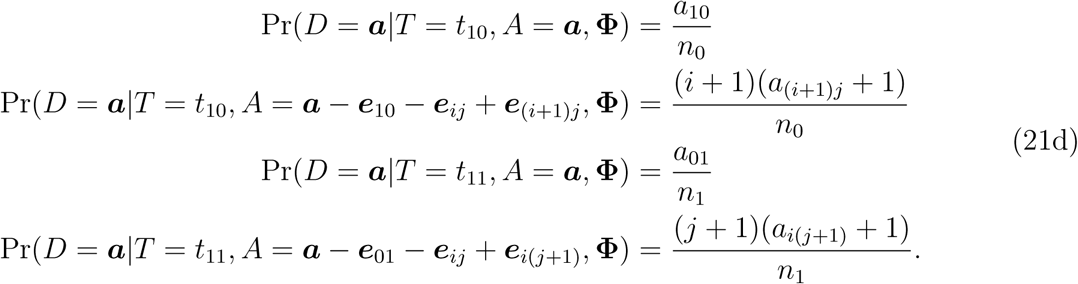

#### 3.2.3 Migration events

Ancestor *A* may also be formed by a migration event (*t*_20_ or *t*_21_). Irrespective of the allelic types of the genes, the probabilities of these events correspond to

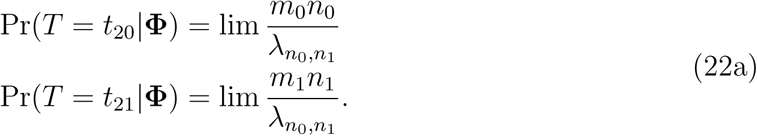

Compatible ancestors correspond to

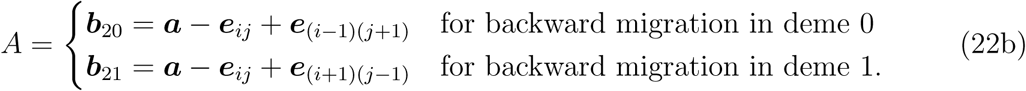

For these ancestors, we have

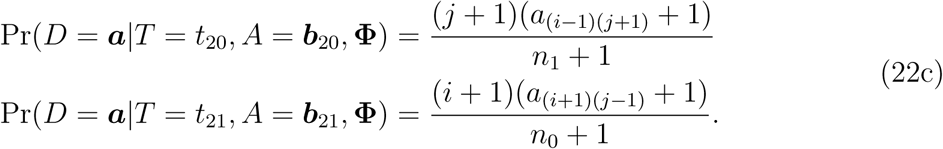

As in the cases of coalescence and mutation, these expressions reflect forward-in-time probabilities of generating descendant *D* from ancestor *A* under selective neutrality.

## 4 Induction

We determine the probabilities of the allele frequency spectra by progressively incrementing sample size (*n*), number of alleles (*K*), and number of singletons (*S*). This inductive approach mirrors the original argument of Ewens (1972).

Our base case corresponds to samples of size *n* = 2. A monomorphic sample (*K* = 1) occurs with probabilities given by ***G*_2_**(0) (15). Otherwise, *K* = 2 alleles are observed, with the probabilities of monomorphism and polymorphism summing to unity for any given sampling configuration (numbers of alleles across demes).

For a given sample size (*n* ≥ 3), we consider successively larger numbers of alleles (*K ≤ n*). For any sample size *n*, evaluation at *z* = 0 of the vector of probability generating functions ***G_n_***(*z*) (14) for the number of alleles in the sample provides probabilities for monomorphic samples (*K* = 1). For each assignment of *n* and *K*, we consider successively larger numbers of singletons (0 ≤ *S* ≤ *K*, with *S* = *K* only for *K* = *n*).

### 4.1 General linear system

Let ***P_xyz_*** denote the vector of allele frequency spectrum (AFS) probabilities for samples of size *n = x* genes that comprise exactly *K = y* alleles, of which *S = z* are singletons. Descendant *D* in (18) corresponds to an element of this vector. Section 3.2 describes the rates of transition from ***P_xyz_*** back in time through a single evolutionary event (coalescence, mutation, or migration). Central recursion (18) specifies a linear system of equations to be solved for ***P_xyz_***.

A backward migration event from any descendant *D* in ***P_xyz_*** implies an ancestor *A* also in ***P_xyz_***.

Coalescence (20) implies an ancestor *A* with the same number of alleles (*y*) and singletons (*z*), but with smaller sample size (*x* − 1). Such ancestors correspond to elements of vector ***P***_(***x***−1)***yz***_, which will have already been obtained in the course of the induction.

For the most recent event corresponding to mutation, (21) lists the ancestors *A* with positive probability of generating descendant *D*. Forward in time, a mutation that occurs in a lineage representing a singleton allele in *A* preserves the allele frequency spectrum (*D = A*), implying that any such ancestor *A* corresponds to an element of ***P_xyz_***. Alternatively, a mutation in a lineage representing a non-singleton allele in *A* implies an ancestor AFS of the same sample size but with one fewer allelic class and one fewer singleton relative to *D*. The corresponding vector of AFS probabilities, ***P_x_***_(***y***−1)(***z***−1)_, will have already been obtained in the induction as well.

Matrix ***J_0_*** presents rates of transition among states within ***P_xyz_***, with off-diagonal elements corresponding to migration events and diagonal elements to mutation events that preserve the ancestor AFS (mutation in a singleton of *A*). Matrix ***J_1_*** presents rates of transition through mutation from states in ***P_xyz_*** backward to states in ***P***_x(***y***−1)(***z***−1)_. Matrix ***J_2_*** presents rates of transition through coalescence from ***P_xyz_*** to ***P***_(***x***−1)***yz***_. Conversion from transition rates to transition probabilities entails division of each row (representing a particular state in ***P_xyz_***) by the corresponding total rate of transition through migration, coalescence, or mutation (19). Diagonal matrix **Λ** presents these total rates along the diagonal in the order implied by vector ***P_xyz_***(see Appendix A).

Central recursion (18) implies a linear system of equations:

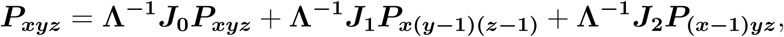

for ***P***_x(***y***−1)(***z***−1)_ and ***P***_(***x***−1)***yz***_ previously determined. Solving, we obtain the AFS probabilities for samples of size *n = x* comprising *K = y* alleles and *S = z* singletons:

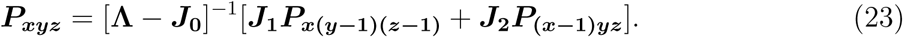

Appendix B presents explicit expressions for a sample of size *n* = 3 genes with *K* = 2 alleles and *S* = 1 singletons.

### 4.2 Horizontal argument

While the vertical (across-time-period) argument which implies (23) can provide all AFS probabilities, we introduce in addition a horizontal (within-time-period) argument that addresses the last gene sampled. This approach facilitates computation of the recursion in likelihoods.

Karlin and McGregor (1972) characterized the last-sampled gene in a sample derived from a panmictic population as a conditional probability of an *n*-gene sample given the penultimate (*n* − 1)-gene sample. We extend this horizontal argument to structured populations to address the probability of spectra that contain at least one singleton allele in deme 0. Analogous expressions hold for AFSs comprising at least one singleton in deme 1.

Starting with AFS ***a*** *−* ***e***_10_, comprising *n*_0_ *−* 1 genes derived from deme 0 and *n*_1_ genes from deme 1, we form the final sample of size *n* = *n*_0_ + *n*_1_ by adding an additional gene from deme 0. The conditional probability that the next-sampled gene represents a novel allele corresponds to

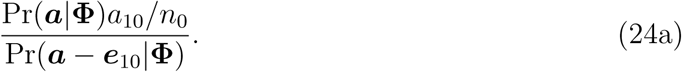

This expression reflects that the addition of a novel allele implies that the AFS of the final sample correspond to ***a***. Because all sampling orders for the formation of ***a*** are equally likely (compare Karlin and McGregor 1972), *a*_10_/*n*_0_ denotes the probability that a singleton in deme 0 is sampled last. Alternatively, we begin with AFS ***a − e***_10_ as before, but add a gene belonging to an allelic class already represented by *i* replicates in deme 0 and *j* replicates in deme 1. In this case, AFS of the sample after the addition of the last gene must correspond to ***a*** *−* ***e***_10_ *−* ***e****_ij_* + ***e***_(*i*+1)*j*_ and the probability that a lineage belonging to such an allelic class is sampled last is (*i* + 1)(*a*_(*i*+1)*j*_ + 1)/*n*_0_. Summing over all possible values of *i* and *j*, we obtain the conditional probability that starting with ***a − e***_10_, the last-sampled gene represents an allele already observed in the sample,

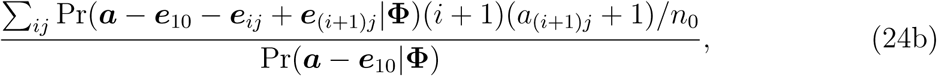

in which unmeaningful expressions (*e.g.*, involving an AFS with negative entries) are defined as zero. These conditional probabilities, that the last-sampled gene represents a novel allele or a non-novel allele, must sum to unity. Accordingly, we may obtain the probability of a sample containing at least one singleton allele by solving for Pr(***a****|***Φ**) in

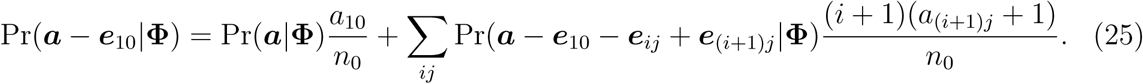

The term on the left corresponds to a sample of smaller size (*n* − 1) than ***a*** and the summation includes spectra of the same sample size (*n*) but fewer singletons. Because our inductive method (Section 4) proceeds by increasing first *n* (sample size) and then *S* (number of singletons) for a given *n*, those two quantities will have already been determined at the point at which Pr(***a***|**Φ**) is addressed. For any allele frequency spectrum ***a*** comprising at least one singleton, recourse to expressions analogous to (25) can simplify the induction described in Section 3.1.

We note that (25) enters in a natural way into the vertical recursion (18). For *t = t*_10_in (18), the most recent event in the genealogical history is a mutation in a lineage in deme 0. The factor of Pr(*T = t*_10_|**Φ**) in (18) corresponds to the right side of (25). The first term on the right represents the generation of descendant *D* = ***a*** from ancestor *A* = ***a*** through mutation in a lineage representing a singleton allele. Proceeding forward in time, this event occurs in *A* with probability *a*_10_/*n*_0_. Alternatively, *D* = ***a*** may be generated through mutation in one of the (*i* + 1) replicates in deme 0 of an allelic class that also has *j* replicates in deme 1 of ancestor *A* = ***a − e***_10_ − ***e****_ij_* + ***e***_(*i*+1)*j*_, with probabilities indicated in the summation.

## 5 Exploration

While the primary purpose of this note is to present our inductive method for determining the probability of allele frequency spectra for samples derived from a structured population, we here indicate some avenues of exploration.

### 5.1 Simulation results

Figure 1 presents an example of the correspondence between simulation results using ms (Hudson 2002) and the AFS probabilities determined inductively from (23). Depicted are only a small subset of the 257 spectra possible for a 10-gene sample comprising *n*_0_ = 3 genes derived from deme 0 and *n*_1_ = 7 from deme 1. Here, we assign *θ* = 1 and assume symmetry in effective numbers and migration rates (9).

**Figure 1:**
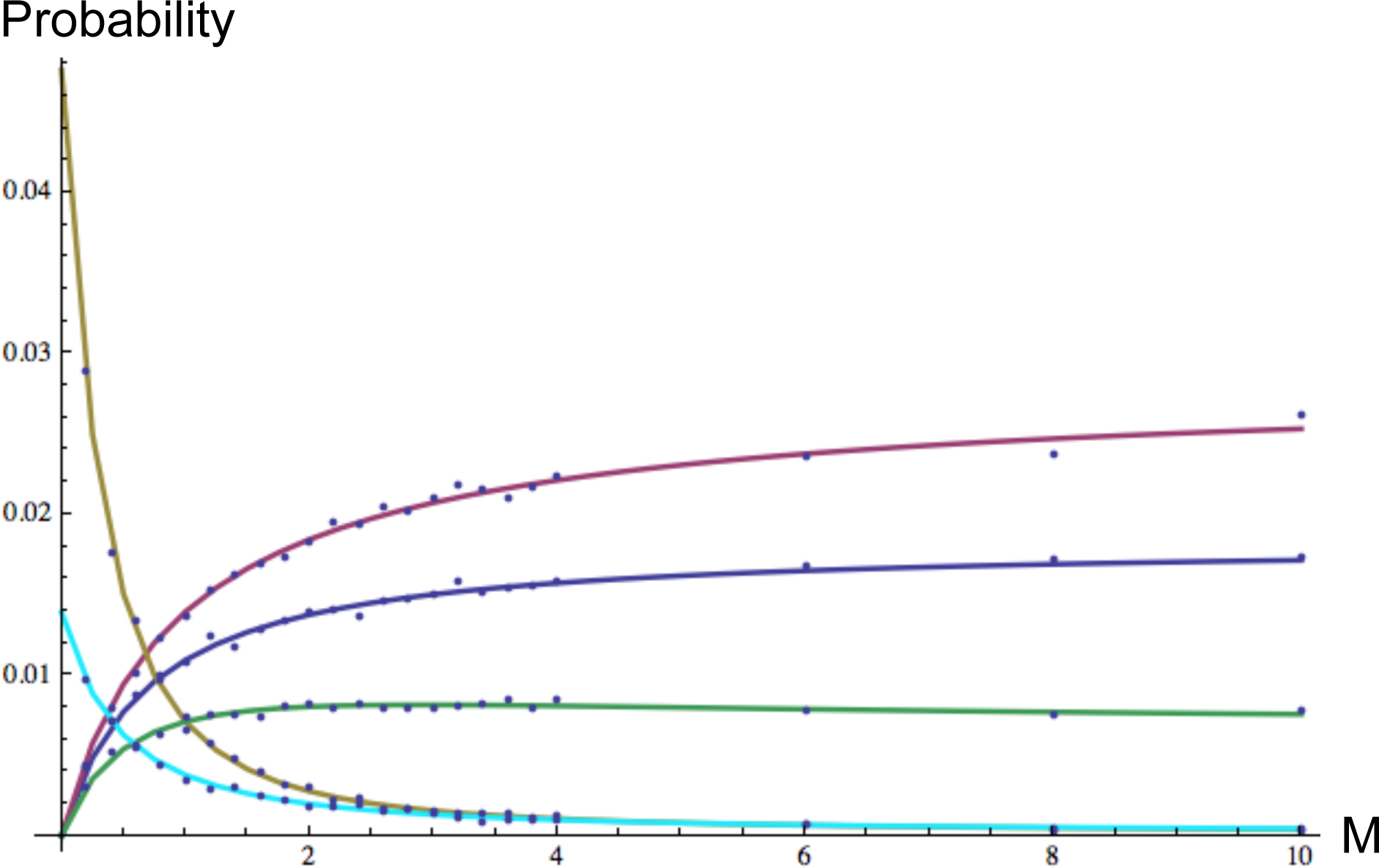
Allele frequency spectrum probabilities for a sample comprising *n*_0_ = 3 genes derived from deme 0 and *n*_1_ = 7 from simulations (dots) using ms (Hudson 2002) and analytical expressions (lines) obtained from deme 1 under symmetry (9) and *θ* = 1, as a function of *M*_0_ = *M*_1_ = *M*. These curves illustrate some of the trends exhibited by the 257 spectra possible for this sampling configuration. Each dot represents the proportion of 10,000 independent simulations generated by ms (Hudson 2002) that correspond to the indicated AFS. To convert the infinite-sites mutation scheme of ms to infinite-allele mutation, we designate an allelic class as a particular haplotype, with distinct alleles differing by at least one substitution. Lines indicate analytical values of spectrum probabilities obtained inductively from (23): ***e***_01 +_ ***e***_36_ (2 alleles, cherry), ***e***_37_ (1 allele, blue), ***e***_01_ + ***e***_03_ + ***e***_33_ (3 alleles, green), ***e***_07_ + ***e***_30_ (2 alleles, sand), and ***e***_03_ + 2***e***_02_ + ***e***_30_ (3 alleles, teal).

Complete isolation presents a discontinuity in the model, with each allele restricted to a single deme. Small but positive rates of backward migration permit many more spectra.

For low rates of migration (small *M*_0_ = *M*_1_ = *M*), the two AFSs shown in Figure 1 with no alleles represented in both demes have higher probability: {***e***_07_ + ***e***_30_} (sand, with *K* = 2 alleles) and ***e***_03_ + 2***e***_02_ + ***e***_30_ (teal, *K* = 4). For larger migration rates, the probabilities of those AFSs declines with increases in the probabilities of AFSs with alleles present in both demes: {***e***_01_ + ***e***_36_} (cherry, *K* = 2), {***e***_37_} (blue, *K* = 1), and {***e***_01_ + ***e***_03_ + ***e***_33_} (green, *K* = 3).

### 5.2 Effective mutation rate approximation

Among the most pervasive implications of population structure on the pattern of genetic variation is its influence on the coalescence process. Just as coalescence cannot be the most recent evolutionary event between lineages belonging to distinct allelic classes, it cannot occur between lineages presently residing in distinct demes. Were it the case that an effective value of *θ*, the sole parameter of the ESF, could accommodate population structure under general assignments of the parameters (6), the ESF together with its remarkable properties would provide an inference framework for the analysis of samples derived from a wide range of ecological contexts. Unfortunately, such an effective *θ* does not exist: even for very small samples, counterexamples are simple to produce that show that the candidate solution obtained for one AFS does not hold for other AFSs. Even so, we address whether an effective scaled mutation rate can provide a useful approximation of salient features of the allele frequency spectra under the relatively simple case of symmetric migration (*M*_0_ = *M*_1_ = *M*).

#### 5.2.1 Candidates

A simple candidate for an effective mutation rate interpolates between the extremes for *M*:

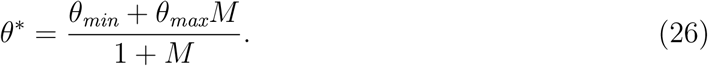

Choices for *θ_min_* include

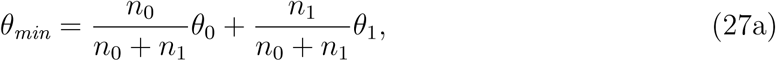

for *θ_i_* = lim 4*N_i_u*, or the solution of

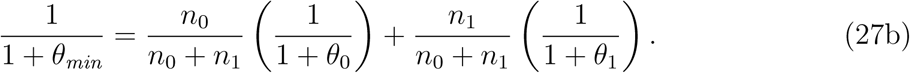

In considering expressions for *θ_max_*, we note that high migration rates not only increase the opportunity for lineages presently in distinct demes to transfer to the same deme, but also increase the rate of separation of lineages presently in the same deme. To capture this effect, we consider the influence of mutation on the probability of identity between a random pair of genes. In the absence of population structure, the ESF (3) provides the probability of a monomorphic sample of size *n* = 2:

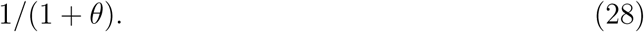

For a structured population, ***G_2_***(0) (15) provides the probability that a sample of size *n* = 2 is monomorphic under the scaling (8). The probability that a random pair, sampled without regard to deme of origin, belong to the same allelic class is

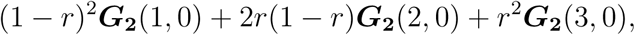

for *r* (5) the proportion of the total effective number of individuals residing in deme 0 and ***G_2_***(*i*, 0) the *i^th^* element of ***G_2_***(0) (15), corresponding to *i* − 1 genes sampled from deme 0. Setting this expression equal to the probability of identity under panmixis (28), we obtain a candidate effective mutation rate *θ^*^* by solving

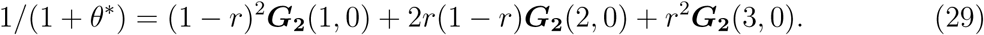

At the low-migration limit (*M*_0_, *M*_1_ *→* 0), (15) implies

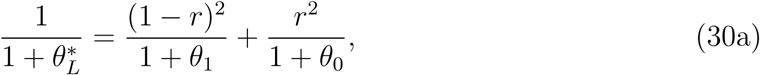

for *θ_i_* = lim 4*N_i_u*. At the high-migration limit (*M*_0_*, M*_1_), all elements of ***G_2_***(0) reduce to (17), implying an effective mutation parameter of

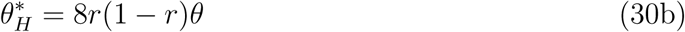

under (8). This expression indicates that at the high-migration limit,

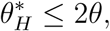

with equality only for demes of equal size (*N*_0_ = *N*_1_, *r* = 1/2). Population structure does not necessarily imply increased effective number; in particular, the high-migration 
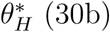
 is less than *θ* for

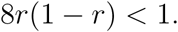

#### 5.2.2 Distance between spectra

We use the Bhattacharrya distance to compare the allele frequency spectrum (AFS) under population structure (23) to an approximation based on the ESF (3) with an interpolated effective *θ* function (26). This index of distance views the spectrum probabilities as a vector with elements summing to unity. In a comparison of a pair of discrete probability distributions,

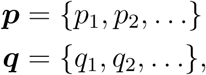

in which the *i^th^* element represents the probability of the *i^th^* spectrum, the vectors are standardized to length 1 by replacing each element by its square root. The Euclidean distance between the standardized vectors corresponds to the Bhattacharrya coefficient. The Bhattacharrya distance represents a further normalization to constrain the coefficient to [0, 1] by taking the natural logarithm:

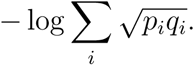

We determine the Bhattacharrya distance between the ESF (3) using an effective *θ* and the actual AFSs (23), collapsed by removing information about the number of replicates of the alleles in each deme. To obtain the collapsed distribution, we assign each full AFS to an equivalence class, defined by a specified number of alleles (*K = k*) and total number of replicates of each allele. For example, the equivalence class characterized by ***c*** = *{c_x_}* (2) includes AFS ***a*** = *{a_ij_}* if

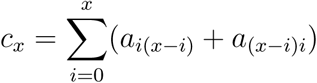

for all *x*. We then determine the difference between the probability of ***c*** under the ESF (3) and the sum of the probabilities of full AFSs that belong to this equivalence class.

Here, we illustrate the departures between the actual AFS and the approximation using the interpolated effective *θ* candidate (26) with (27a) substituted for *θ_min_* and (30b) for *θ_max_*. Figure 2 presents the Bhattacharrya distance between the collapsed AFS and the approximation for a sample comprising *n* = 10 genes, with *n*_0_ derived from deme 0 and *n*_1_ from deme 1. We set the proportion of the total population that resides in deme 0 to *r* = 0.3 (7) and the actual scaled mutation rate *θ* = lim 2(*N*_0_ + *N*_1_)*u* to unity. We present 4 sampling configurations: all genes derived from deme 0 (*n*_0_ = 10, green), *n*_0_ = 5 (red), *n*_0_ = 3 (blue), and *n*_0_ = 0 (black). For all configurations, the distance between the approximation and the collapsed AFS becomes small for very high migration rates (*M*).

**Figure 2:**
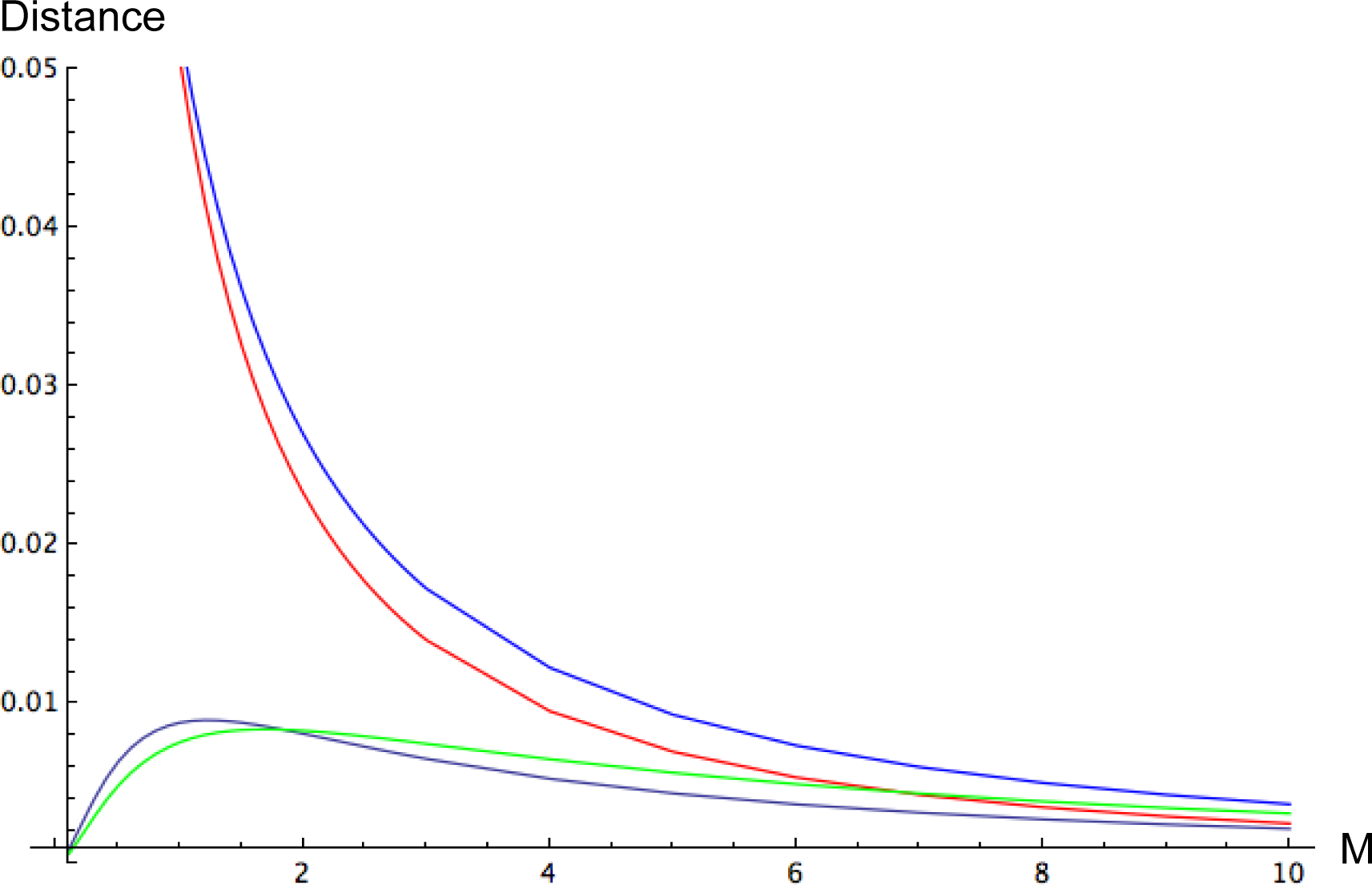
Bhattacharrya distance between the actual allele frequency spectrum (23), collapsed to remove information about location of lineages, and the ESF (3) using effective mutation function (26) with *θ_min_* corresponding to (27a) and *θ_max_* to (30b). The full AFS probabilities reflect a sample size of *n* = 10 genes, of which *n_i_* derive from deme *i* (*i* = 0, 1), the residence in deme 0 of a proportion *r* = 0.3 of the total population (7), scaled backward migration rate *M*_0_ = *M*_1_ = *M*, and *θ* = lim 2(*N*_0_ + *N*_1_)*u* = 1. The green line corresponds to a sample derived entirely from deme 0 (*n*_0_ = 10, *n*_1_ = 0), red to *n*_0_ = 5, blue to *n*_0_ = 3, and black to *n*_0_ = 0.

Very low migration rates (*M* → 0) promote the more recent occurrence of coalescence events compared to migration events. In such cases, spectra in samples derived entirely from a single deme (green and black) converge as expected to the panmictic ESF (3) with the local mutation parameter (*θ_i_*). As *M* increases, the discrepancy between the actual and approximate AFS grows, reaching an intermediate maximum before declining for very large *M*. The discrepancy persists for larger *M* in the sample derived entirely from the smaller deme (green line).

Sampling configurations that comprise genes from both demes (red and blue) show large departures between the actual and approximate AFSs even for substantial values of *M*. For very low migration rates (*M* → 0), the actual AFS converges to a mixture of ESF distributions with distinct *θ* parameters rather than to a single ESF distribution with an interpolated parameter (26). Convergence to the panmictic approximation appears to require migration rates rather beyond the “one-migrant-per-generation” rule used in many conservation studies, which in our analysis corresponds to values of *M_i_* = 4*N m_i_* around 4 or less, depending on the definition of the scaling factor *N* (6).

#### 5.3 Site frequency spectrum

As described in Ganapathy and Uyenoyama (2009), we derive the the folded site frequency spectrum (SFS) by conditioning the the ESF (3) on biallelic samples (*K* = 2), using (14). We note that this approach yields the probability mass function for the SFS under arbitrary mutation rates, not only expectations of the number of derived alleles in a sample under the assumption of a vanishingly small mutation rate or limitation to a single mutation in the sample genealogy (*e.g.*, Fu 1995; Hudson 2015).

The striking finding of Ewens (1972) that the ESF (3) conditioned on the number of alleles in the sample (*K*) is independent of the mutation parameter *θ* no longer holds under population structure. While a fuller treatment of the effects of population structure on the SFS will appear separately, we here illustrate some properties for a sample of size *n* = 10 derived from a structured population under symmetry (9).

Figure 3 presents a comparison of the distribution of the folded SFS expected under panmixis to the actual distribution under population structure (23) with mutation parameter assigned as *θ* = 0.1. Under low migration (*M* = 0.1, red), the actual SFS (red) corresponds relatively well to the panmictic SFS (black) for samples derived entirely from a single deme (*n*_0_ = 0). For all other sampling configurations, the probability of the partition that mirrors the distribution of genes across demes exceeds the probability under panmixis. Low migration rates promote rapid coalescence among genes sampled from the same deme relative to migration events. This process tends to favor gene genealogies in which a long branch partitions the sample by deme of origin. The occurrence of the mutation that generates the biallelic sample on that long branch would induce the observed pattern. Substantial departures from the panmictic case (black) persist even for migration rates on the order of one migrant per generation (*M* = 1.0, blue).

**Figure 3:**
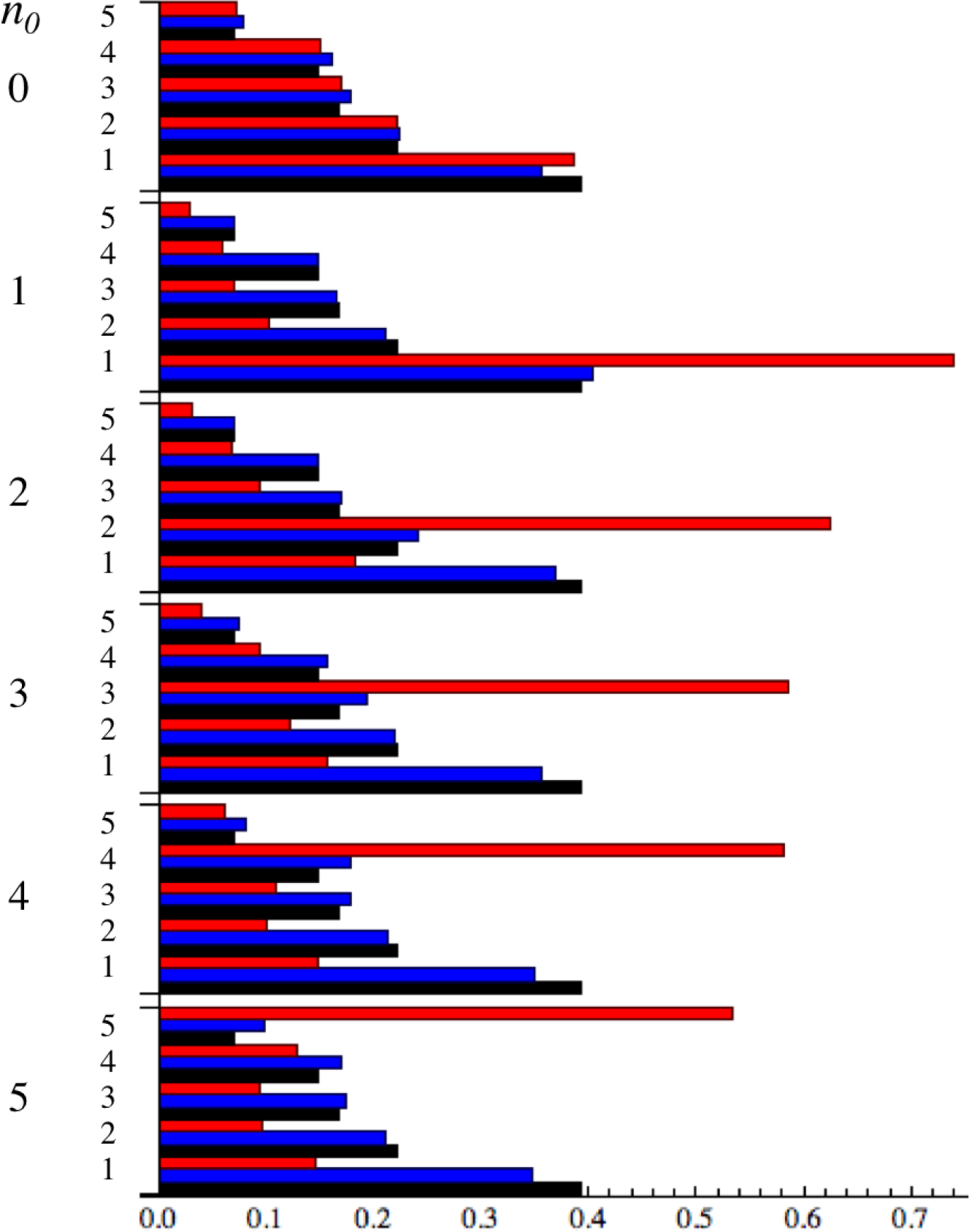
Folded site frequency spectra (SFSs) for a sample comprising *n* = 10 genes under symmetry (9), with *M*_0_ = *M*_1_ = 0.1 and *θ* = 0.1. Listed under the column heading *n*_0_ is the number of genes derived from deme 0, with the complement (*n*_1_ = 10 − *n*_0_) derived from deme 1. The vertical scale indicates the number of observations of the minor allele in the sample (without distinguishing between the derived and ancestral alleles). Histograms depict allele frequency spectra conditioned on *K* = 2 alleles, including the actual spectrum probabilities (23) under *M* = 0.1 (red) and *M* = 1.0 (blue) compared to the ESF (3) (black).

Figure 4 compares departures of the actual folded SFS obtained from (23) to the panmictic expectation (black) under even higher rates of migration (red: *M* = 2, blue: *M* *=* 4) for two mutation intensities (*θ* = 1, 5). For *θ* = 1 (left), very high migration (*M* = 4, blue) appears to reduce the departure of the actual SFS from the panmictic expectation (black), although a deficiency in the singleton class relative to the panmictic case persists. However, higher mutation (*θ* = 5, left) intensifies the pattern shown in Fig. 3: an excess of the class that conforms to the initial partition of the sample between demes (*n*_0_). This trend illustrates that in the full model, reflecting coalescence, mutation, and, migration, the effect of any given process depends on its rate relative to the other processes.

**Figure 4:**
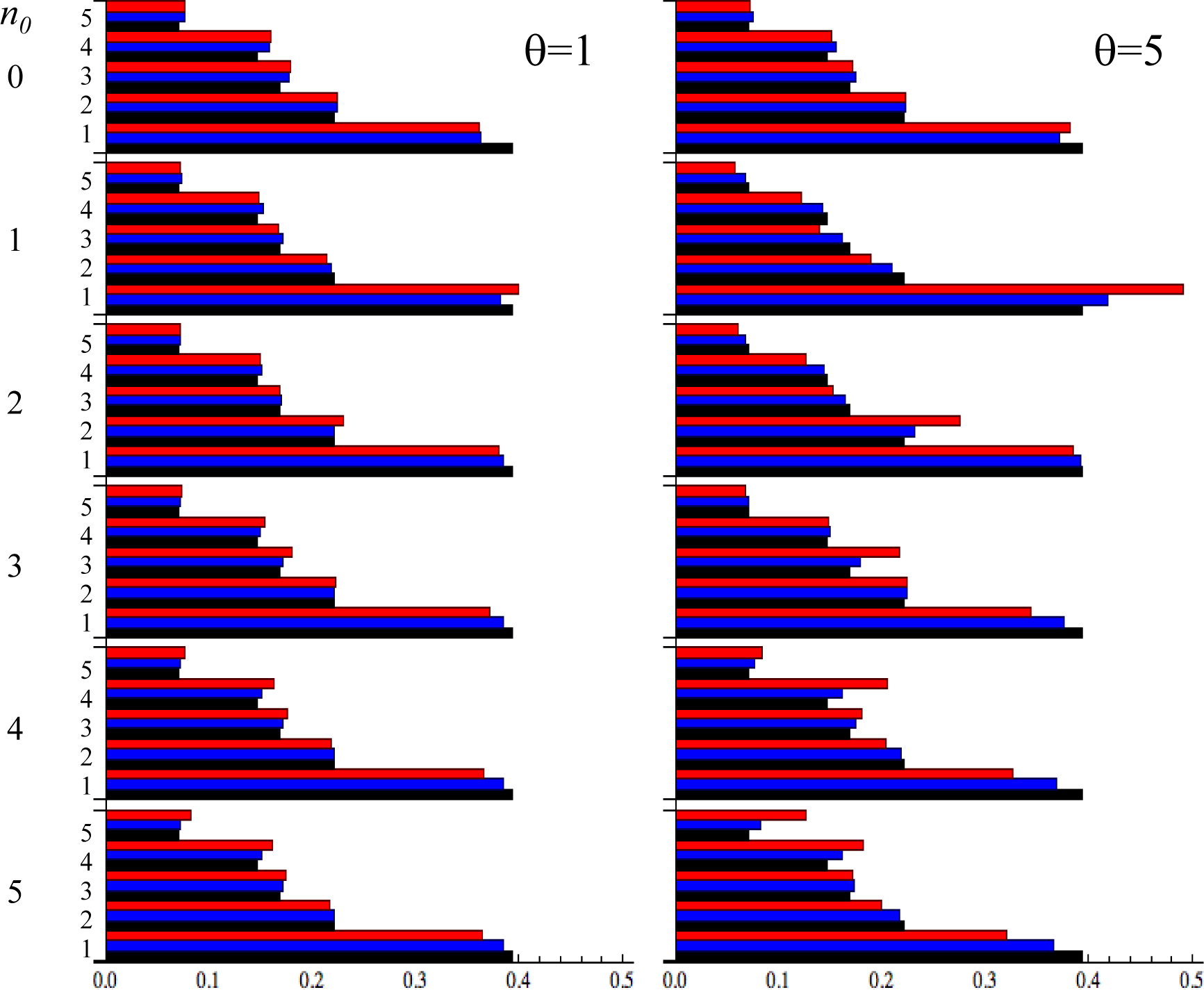
Folded site frequency spectra (SFSs) for a sample comprising *n* = 10 genes under symmetry (9), with *θ* = 1 (left) and *θ* = 5 (right), with axes similar to Fig. 3. Histograms depict allele frequency spectra conditioned on *K* = 2 alleles, including the ESF (3) (black) and actual spectrum probabilities (23) for *M* = 2 (red) and *M* = 4 (blue).

## 6 Discussion

Our method (23) inductively determines exact allele frequency spectrum (AFS) probabilities under the infinite-allele mutation model for a sample derived from a population comprising two demes. This approach also provides probability generating functions (14) for the number of alleles observed in a sample of a specified total size and configuration across demes.

### 6.1 Allele frequency spectra in structured populations

Our analysis addresses a class of structured population models for which the importance sampling (IS) methods of De Iorio and Griffiths and colleagues (*e.g.*, De Iorio and Griffiths 2004; De Iorio *et al.* 2005) can generate approximations to the likelihood. Our method addresses only infinite-allele mutation, while their method is designed to accommodate other mutation models as well. Our preliminary comparisons to the exact AFS probabilities suggest that their IS proposals are indeed efficient. Even so, determination of likelihoods without resort to intensive sampling may serve to improve the feasibility of Bayesian and other approaches in practical applications to genetic data.

As backward migration rates become arbitrarily large, population structure declines, with AFS probabilities converging to the ESF (Fig. 1). However, the level of migration required for a close correspondence of the AFS to the panmictic case appears to lie beyond the “one-migrant-per-generation” rule for many sampling configurations (Fig. 2).

An immediate result from an examination of spectrum probabilities for even very small samples is that population structure cannot in general be accommodated by preserving the ESF (3) but substituting an effective value of *θ*, its sole parameter. Even so, we have explored whether a qualitative portrait of the AFS can be generated by defining an effective *θ* (26) as a simple interpolation between complete isolation (*M* = 0) and near panmixia (*M* → ∞). This approximation appears to provide some guidance for samples derived entirely from a single deme, but less so for other sampling configurations (Fig. 2). In practice, empiricists may be unaware of the nature of structure in the population from which a sample is derived, much less the numbers of genes derived from each deme. Accordingly, prospects for accommodating population structure simply by adopting an effective *θ* value appear dim.

### 6.2 Effects of substitution rate on site frequency spectra

Among the key properties of the ESF (3) is that the allele frequency spectrum (AFS) conditioned on the number of alleles (*K*) in the sample is independent of the scaled mutation rate *θ* (Ewens 1972). For panmictic populations, Ganapathy and Uyenoyama (2009) demonstrated that conditioning on a single mutational event in the history of the sample introduces distinct constraints, with very high *θ* values tending to generate an excess of rare alleles relative to the ESF conditioned on *K* = 2 alleles. We find that under population structure, *θ* affects the shape of the AFS even after conditioning on the number of alleles in the sample. In particular, population structure affects the folded biallelic site frequency spectrum (SFS), which corresponds to the AFS conditioned on *K* = 2 alleles (Ganapathy and Uyenoyama 2009).

Analyses of genome-wide patterns of variation in the Exome Aggregation Consortium (ExAC) data set have revealed the strong influence of mutation rate on the SFS (Harpak *et al.* 2016; Lek *et al.* 2016). Recurrent mutation and selection may account for a substantial part of this departure from neutral expectation under panmixis. Our study suggests that population structure alone is expected to induce a dependence of the SFS on the rate of neutral substitution. Human population structure may be complex, particularly in the planet-wide data set assembled by the ExAC.

A widely-used approach to controlling for demographic history entails comparing patterns of variation among functional classes (*e.g.*, Kim *et al.* 2007). Under population structure, however, the substantial variation in *θ* among functional classes (Li *et al.* 1981; Bustamante *et al.* 2003) may also contribute to variation in SFS shape across classes. Further, the rapid and pervasive changes in mutation rates through time that have occurred in recent human history (Harris and Pritchard 2017) may also affect inferences about departures from neutrality derived from SFS analyses.

## Acknowledgments

We thank Marcus W. Feldman for the extraordinary inspiration and support that has sustained us throughout our careers. We are grateful to Sohini Ramachandran and Noah Rosenberg for the opportunity to contribute to this volume honoring Marc and for their patience and support. We thank Benjamin D. Redelings for his help in an early phase of development. Public Health Service grant GM 37841 (MKU) provided partial funding for this research.

## Appendix A Markov chain description

We characterize the system as a finite-state, continuous-time Markov chain. For a given level of the gene genealogy, the Markov chain describes the process proceeding backward in time, from the entrance state (most recent) to the exit state (passage to the preceding level).

### A.1 Transition rates

Let ***P*** (*t*) denote the transition probability matrix, of which the *ij^th^* element represents the probability that a process exists in state *j* at time *t*+*s*, given its residence in state *i* at time 0. With respect to entrance state *i*, we denote as transient any state *j* for which 
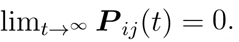
. Termination of the level corresponds to absorption in exit state *j* (***P*** *_jj_* (*t*) = 1 for all *t*). We restrict attention to processes in which all configurations can be classified as either transient or absorbing.

Under the Markov properties, ***P*** (*t*) satisfies the Chapman-Kolmogorov equations:

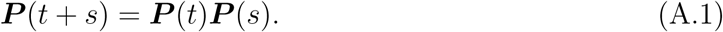

In particular,

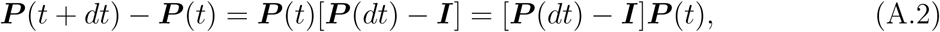

for ***I*** the identity matrix and *dt* a small time increment, with instantaneous rates of change given by

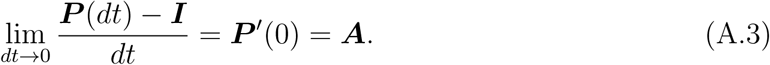

From (A.2) we obtain

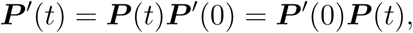

the solution of which gives

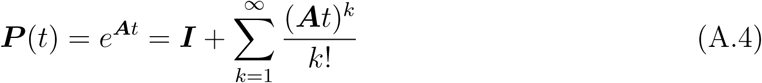

(see, for example, Taylor and Karlin 1998, Chapter VI, section 6).

Beginning from the entrance state to a genealogical level, the process may visit some number of transient states before absorption in an exit state (on the next level back in time). For the two-deme model under consideration, transitions among transient states correspond to backward migration of lineages among demes and transitions to exit states to coalescence of a pair of lineages residing in the same deme. For *x* the number of transient states accessible from the entrance state and *y* the total number of exit states accessible from any of the transient states, the instantaneous rates of change (A.3) correspond to

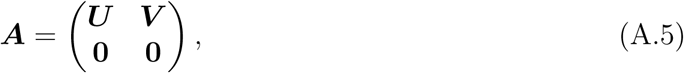

in which ***U*** (dimension *x × x*) gives the instantaneous rates of within-level moves and ***V*** (*x × y*) between-level moves, with the lower blocks representing matrices of zeroes of appropriate size (*y × x* and *y × y*). From (A.4) and (A.5), we obtain the transition probability matrix

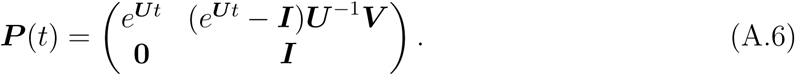

### A.2 Event probabilities

An alternative formulation characterizes the coalescence process in terms of probabilities (rather than rates) of transitions from the present state (see Leman *et al.* 2005; Kumagai and Uyenoyama 2015). Let *λ*(*i*) represent the total instantaneous rate of change from transient state *i*, the sum of the off-diagonal elements of row *i* of ***A***, with **Λ** a diagonal matrix of the *λ*(*i*). The diagonal elements of ***U*** correspond to **Λ**. Matrices of probabilities (rather than rates) of transitions correspond to

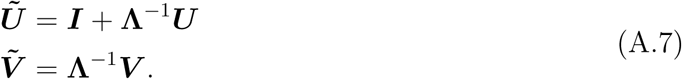

For a process initiated at transient state *i*, the distribution of the states after *z* transitions to a transient state corresponds to the *i^th^* row of ***Ũ*** *^z^*. Let ***b*** denote the transient state from which the transition to an absorbing state occurs following any number of transitions among transient states (*z* = 0, 1*, …,*). For a process initiated at state ***a***, the distribution of absorption from state ***b*** corresponds to the *ij^th^* element of

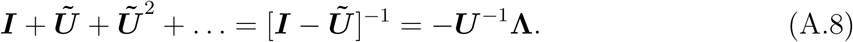

for *i* the index of state ***a*** and *j* the index of state ***b***.

### A.3 Hitting probabilities

That ***U*** in (A.6) represents rates of transition among transient states implies

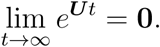

Accordingly, the probability that a process presently in transient state *i* exits through state *j* corresponds to the *ij^th^* element of

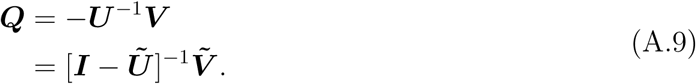

## Appendix B Explicit determination of AFS probabilities for a 3-gene sample

We provide explicit expressions for the components of (23), for the determination of the probabilities of spectra comprising *n* = 3 genes, *K* = 2 alleles, and *S* = 1 singletons.

A total of six spectra of this kind exist, with probabilities given by

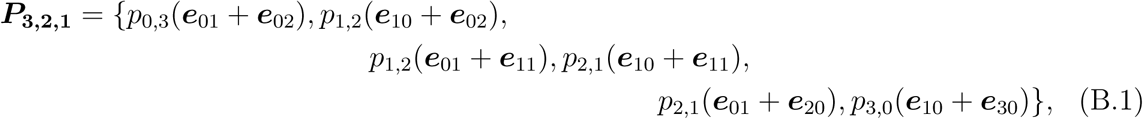

in which *p_ij_* denotes a sample comprising *i* genes from deme 0 and *j* genes from deme 1 and ***e_kl_*** a unit vector with the *kl* element corresponding to 1 and all other elements to 0.

Coalescence can occur only between a pair of lineages that both reside in the same deme and represent the same allelic class. In this 3-gene sample, non-singleton alleles correspond to doubletons (***e***_02_ or ***e***_20_). Accordingly, coalescence implies an ancestor with reduced sample size (*n* = 2), the same number of alleles (*k* = 2), and an additional singleton (*S* = 2):

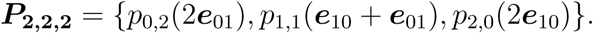

Mutation events imply an ancestor with the same sample size (*n =* 3) that either are identical to the descendant or comprise one fewer singleton:

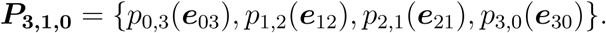

For the ordering of states indicated in (B.1), the total rates of change (19) correspond to

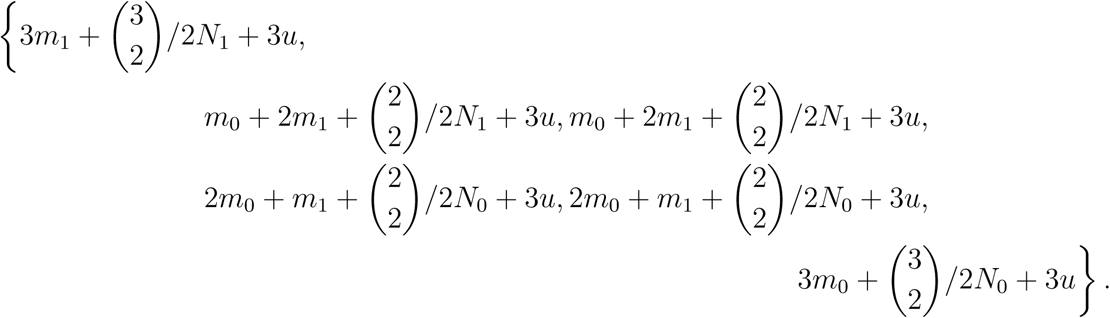

This vector appears on the diagonal of diagonal matrix **Λ**.The other matrices in (23) correspond to

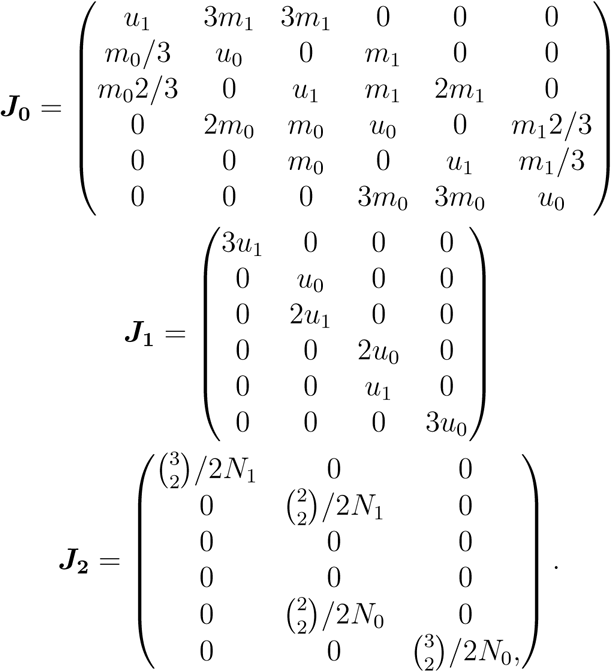

